# Impaired miRNA processing by DICER1 downregulation endows thyroid cancer with increased aggressiveness

**DOI:** 10.1101/373746

**Authors:** Julia Ramírez-Moya, León Wert-Lamas, Garcilaso Riesco-Eizaguirre, Pilar Santisteban

**Affiliations:** Instituto de Investigaciones Biomédicas “Alberto Sols”; Consejo Superior de Investigaciones Científicas (CSIC), Universidad Autónoma de Madrid (UAM), Madrid, Spain.; Departamento de Endocrinología y Nutrición, Hospital Universitario de Móstoles, Madrid, Spain.; Centro de Investigación Biomédica en Red de Cáncer (CIBERONC) Instituto de Salud Carlos III (ISCIII), Madrid, Spain.

**Keywords:** DICER1, microRNAs, miR146b, Enoxacin, Thyroid cancer

## Abstract

The global downregulation of miRNAs (miRs) is emerging as a common hallmark of cancer. However, the mechanisms underlying this phenomenon are not well known. We identified that the oncogenic miR-146b-5p attenuates miRNA biosynthesis by targeting DICER1 and reducing its expression. DICER1 overexpression inhibited all the miR-146b-induced aggressive phenotypes in thyroid cells. Systemic injection of an antimiR-146b in mice with orthotopic thyroid tumors suppressed tumor growth and recovered DICER1 levels. Notably, DICER1 downregulation promoted proliferation, migration, invasion and epithelial-mesenchymal transition through miR downregulation. Our analysis of TCGA revealed a general decrease in DICER1 expression in thyroid cancer that was associated with a worse clinical outcome. Administration of the small molecule enoxacin to promote DICER1 complex activity reduced tumor aggressiveness both *in vitro* and *in vivo*. Overall, our data establish DICER1 as a tumor suppressor and that the oncogenic miR-146b contributes to its downregulation. Moreover, this study highlights a potential therapeutic application of RNA-based therapies including miR-inhibitors and restoration of the biogenesis machinery, which may provide treatments in for thyroid and other cancers.

## Introduction

MicroRNAs (miRs) are short noncoding RNAs that function post-transcriptionally to suppress gene expression through interactions of their seed region with complementary sequences in the 3′-untranslated regions (UTRs) of target messenger RNAs, which consequently affects myriad cellular and developmental pathways. Because a single miR can target different mRNAs, and individual mRNAs can be coordinately suppressed by several miRs, the miR biogenesis pathway has an important role in gene regulatory circuits (Gregory & Shiekhattar, 2005;Lin & Gregory, 2015). Recently, miRs have taken center stage in molecular oncology due to their participation in the development and progression of human neoplasms, including thyroid tumors (Pallante et al, 2014; Riesco-Eizaguirre & Santisteban, 2016).

Thyroid cancer is the most frequent endocrine malignancy and its incidence is the most rapidly increasing among all cancers in the United States (Lim et al, 2017). The classical view of thyroid cancer pathogenesis considers thyroid carcinomas as tumors accumulating mutations that drive progression through a dedifferentiation process initially giving rise to the well-differentiated carcinomas – papillary (PTC) and follicular (FTC) – and progressing to poorly differentiated (PDTC) and undifferentiated or anaplastic (ATC) thyroid carcinomas (Haugen & Sherman, 2013).

Activation of oncogenes is a known cause of miR deregulation in thyroid cells. Among the most well-studied oncogenes that drive malignancy in thyroid tumors are BRAF, RAS and RET/PTC, which induce a distinct set of global changes in the expression of miRs (Fuziwara & Kimura, 2017). Genomic analysis of PTC tumors in The Cancer Genome Atlas (TCGA) suggests that miR expression patterns define clinically relevant subclasses and may contribute to loss of differentiation and tumor progression (Cancer Genome Atlas Research Network, 2014; Riesco-Eizaguirre & Santisteban, 2016). Indeed, such an analysis highlighted six miR clusters, and two of them, clusters 5 and 6, were associated with less-differentiated tumors and higher risk of recurrence. Moreover cluster 5 was enriched for high levels of miR-146b, miR-375, miR-221 and miR-222, and low levels of miR-204, whereas cluster 6 was characterized by high levels of miR-21, miR-221, miR-222, and low levels of miR-204 ( Cancer Genome Atlas Research Network, 2014; Riesco-Eizaguirre & Santisteban, 2016).

Several studies have shown that miR expression is globally suppressed in tumor cells (Lin & Gregory, 2015; Lu et al, 2005; Martello et al, 2010; Thomson et al, 2006). However, little is known about the underlying mechanisms and the phenotypic advantages provided to cells by reduced miR expression. Interestingly, in thyroid cancer, differentiated PTC and FTC present both up- and down-regulated miRs, whereas dedifferentiated and aggressive ATC show almost exclusively down-regulated miRs, with various reports describing 44 down-regulated miRNAs and only 6 up-regulated miRNAs (Braun et al, 2010; Hebrant et al, 2014; Saiselet et al, 2016). This suggests that the progression of differentiated thyroid carcinomas to aggressive ATC is characterized by dramatic changes in miR expression, and that miR downregulation might play a role in this transition. Although the mechanism by which miRs are under-expressed in cancer remains unknown, it could involve the RNAse III-type enzyme DICER1, which plays a fundamental role in processing miR precursors into mature miRs (Gregory & Shiekhattar, 2005). In the canonical miR biogenesis pathway, DICER1 associates with transactivation-responsive RNA-binding protein (TRBP), forming the DICER-complex, to ultimately bind a mature miR strand to the RISC-complex to complementary target mRNAs. In general, lower levels of DICER1 have been found to correlate with worse outcomes in lung, breast, skin, endometrial and ovarian cancer (Foulkes et al, 2014). Yet, no components of the miR-processing pathway have been reported to be completely nonfunctional in human tumors, which is not surprising given that germline deletion of *Dicer1* in mice is lethal (Bernstein et al, 2003). By contrast, recurrent somatic mutations have been detected in metal-binding residues within the RNase IIIb domain of DICER1, leading to a global loss of 5p-derived miRs and decreasing the expression of some tumor-suppressive miRs, which likely helps to explain the selective pressures that give rise to this specific spectrum of mutations in several cancer types (Lin & Gregory, 2015), including thyroid cancer ( Cancer Genome Atlas Research Network, 2014; Khan et al, 2017; Rutter et al, 2016). Some germline mutations in DICER1 are also frequently found in different types of inherited tumors, leading to the abnormal expression of miRs, termed “DICER1 syndrome”, which courses with a high prevalence of multinodular goiter and an increased predisposition to some cancers, such as differentiated thyroid carcinoma (PTC or FTC) (de Kock et al, 2014; Oue et al, 2008; Rome et al, 2008; Rutter et al, 2016). However, the precise role that DICER1 plays in thyroid tumor progression remains unclear.

It is widely accepted that normal developmental processes and cancer share multiple pathways related to cell proliferation and differentiation. In this respect, conditional deletion of *Dicer1* in thyroid follicular cells during mouse development resulted in a strong reduction in the expression of differentiation markers (Frezzetti et al, 2011; Rodriguez et al, 2012). Furthermore, mice were hypothyroid at birth, and presented characteristics of neoplastic alterations in the thyroid tissue at adult stages (Rodriguez et al, 2012). These findings suggest that miRs are necessary to maintain thyroid tissue homeostasis.

Despite their impact on cancer biology, miR-based cancer therapy is still in its early stages, and almost no studies have addressed thyroid cancer. We recently described that intratumoral administration of a synthetic anti-miR inhibiting miR-146b blocked tumor growth of human thyroid tumor xenografts (Ramirez-Moya et al, 2018). Strategies to restore global miR expression would be a welcome addition to the current therapeutic arsenal for thyroid cancer. In this respect, the small-molecule enoxacin, which promotes miR processing in a TRBP-dependent manner (Shan et al, 2008), was shown to have cancer growth inhibitory effects in a panel of twelve cancer cell lines from seven common malignancies (Melo et al, 2011), although none of these studies were performed in thyroid cancer.

With this information at hand, the objective of our present work was to study the function and regulation of DICER1 in thyroid cancer, and to test whether its impaired function resulted in global miR downregulation contributing to a more aggressive phenotype. Using bioinformatic analysis in human tumors, *in vitro* culture and thyroid tumor mouse models, we demonstrate that one of the mechanisms by which DICER1 is suppressed in thyroid cancer is through the main upregulated miRs, with miR-146b being particularly important. Using loss- and gain-of-function approaches, we provide the first evidence that DICER1 is a tumor suppressor in thyroid cancer, and recovery of DICER1 or miR processing *in vivo* is a beneficial antitumoral approach.

## Results

### he major upregulated miRs in thyroid cancer, including miR-146-5p, target DICER1, promoting in vitro cell proliferation, migration and invasion

Recently, a specific miR signature of highly differentially expressed miRs has been established in thyroid cancer by TCGA ( Cancer Genome Atlas Research Network, 2014) and in a cohort of patients by our laboratory (Riesco-Eizaguirre et al, 2015). Among the top overexpressed miRs in our study were miR-146b-5p, miR-146b-3p, miR-221-3p and miR-222-3p, miR-21-5p and miR-21-3p, and miR-182-5p. To evaluate the functional relevance of this signature for thyroid cancer outcome, we examined the downstream critical targets of those miRs using the *in silico* target screening algorithm miRanda. The results of this analysis identified *DICER1* as a putatively shared target of these main miRs, potentially forming an miR biogenesis regulatory network. Coincidently, the 3’UTR of *DICER1* contained several predicted binding sites for all these miRs, with high probability miRSVR scores (Supplementary Figure 1A). By contrast, almost none of the previously described underexpressed miRs in thyroid cancer, such as miR-204, miR-30a and miR-100 (Supplementary Figure 1A) (Riesco-Eizaguirre et al, 2015), were predicted by miRanda to target *DICER1*. These findings raise the possibility that some upregulated mature miRs act in concert as negative feedback regulators to control *DICER1* expression in thyroid cancer, whereas the downregulated miRs may be indirectly affected by this regulatory loop. In agreement with this, analysis of TCGA database using the Cancer Regulome explorer web tool showed that the expression of the most highly upregulated miRs in PTC – miR-146b-5p, miR-146b-3p, miR-21-3p, miR-21-5p, miR-221-3p and miR-222-3p – negatively correlated with *DICER1* mRNA levels (Supplementary Figure 1A). To question whether those miRs of interest directly target *DICER1* in cells, we performed a focused small-scale screen by transiently transfecting each of these miRs individually into the thyroid cell line Nthy-ori 3-1, which experimentally validated that DICER1 protein levels were reduced in each case (Supplementary Figure 1B).

We previously showed that miR-146-5b is the top overexpressed miR in PTC (Riesco-Eizaguirre et al, 2015) with >35-fold higher expression over normal thyroid tissue and also with a high percentage in the PTC miRome, yielding a reasonable mirSVR score (−0.1471, position nt 1767) (Supplementary Figure 1C) towards DICER1 3’UTR and a good negative correlation of −0.39 to DICER1 expression in TCGA dataset (Supplementary Figure 1D). Moreover, miR-146b is one of the most extensively studied miRs in this tumor type (Riesco-Eizaguirre et al, 2015) and appears to be a prognostic factor for PTC, as it associates with aggressive clinicopathological features and poor clinical outcome (Chou et al, 2013). Some critical targets of miR-146b have been identified in PTC, such as PTEN or CDH1 (Ramirez-Moya et al, 2018); however, the full impact and relevance of this particular miR as an oncogenic player in PTC has not been explored in depth. Because miR-146-5p seems prominent among the miRs studied, and might be important in thyroid cancer, we focused our attention on this miR for our further studies and next evaluated which downstream targets may explain its oncogenic features.

We first corroborated that DICER1 is a *bona fide* target of miR-146b by analyzing its expression in normal human thyroid follicular Nthy-ori 3-1 cells stably overexpressing miR-146b (Nthy-ori-146b) or a control vector (Nthy-ori-Null). Results showed that DICER1 mRNA and protein levels were reduced in cells overexpressing miR-146b (Figure 1A), consistent with the findings of our focused/small-scale DICER1 miR target screen (Supplementary Figure 1B). Moreover, ectopic miR-146b expression significantly reduced the activity of a co-transfected luciferase reporter construct containing the putative miR-146b binding region in the *DICER1* 3’UTR (Figure 1B), confirming direct targeting.

**Figure 1.**
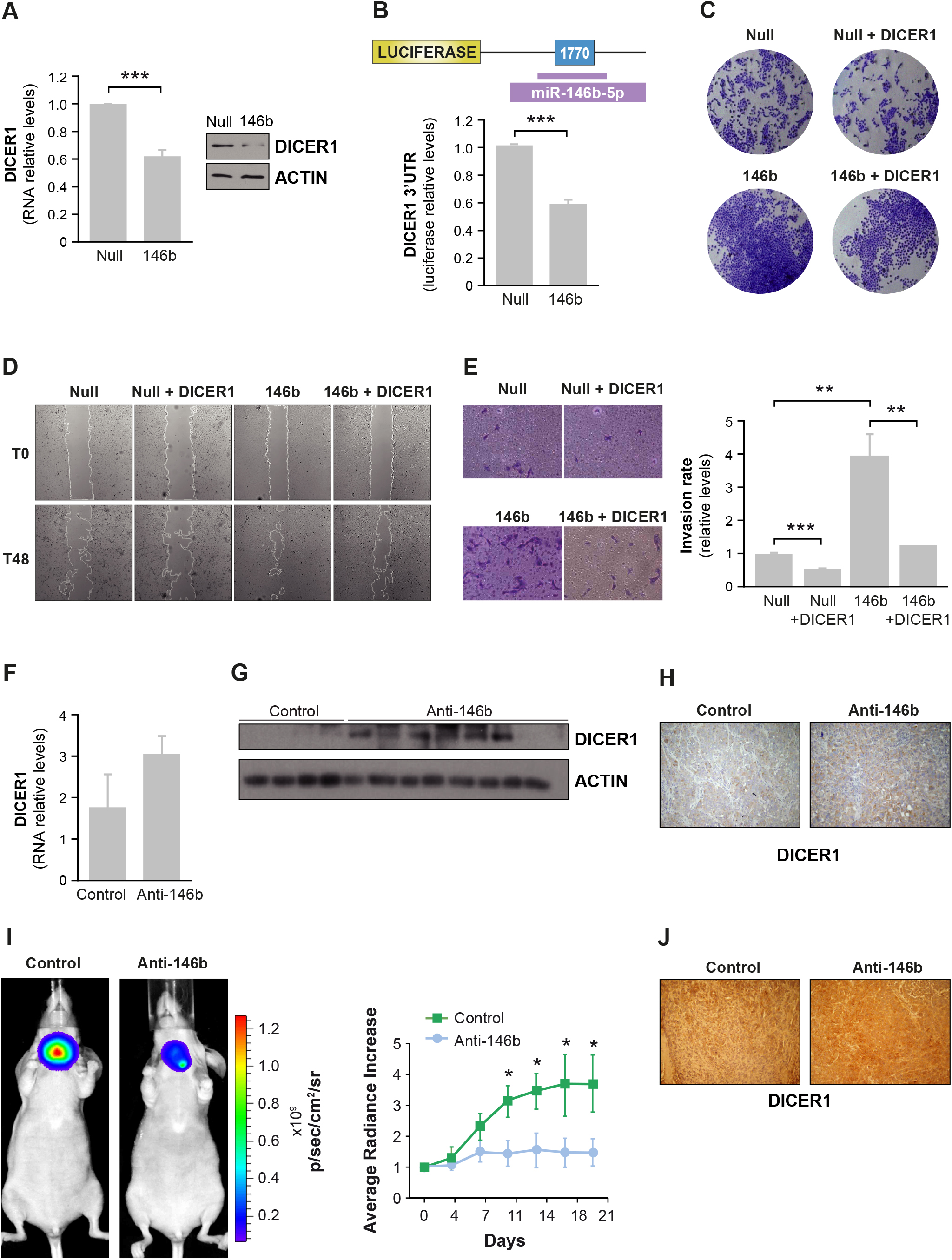
Function of DICER1 and miR-146b-targeted DICER1 *in vitro* and *in vivo* in thryoid cancer establishment. **(A** and **B)** Stable cell lines were generated from Nthy-ori cells transfected with a pEGP-Null vector (Null cells) or a pEGP-miR-146b vector (146b cells). **(A)** Left: relative *DICER1* expression by qPCR. Right: immunoblot of DICER1 expression (results are representative of 3 experiments). **(B)** Direct targeting of DICER1 3´UTR by miR-146b. Luciferase reporter activity relative to *Renilla* level was evaluated in cells 72 h after transfection of pIS1 DICER1 long UTR. **(C)** Representative images of crystal violet-stained Nthy-ori cell lines 48 h after transfection with the DICER1 expression vector. **(D)** Representative images of a wound healing assay 0 and 48 h after scratching. **(E)** Relative quantification of the invasive capacity of cells was analyzed using Matrigel-coated transwell assays. Left: representative images of the lower chamber (invading cells). Right: cell invasion relative to that of “Null” cells. **(F–H)** Tumor samples taken from mice treated with a control or an miR-146b inhibitor (Anti-146b) administered intratumorally (Ramirez-Moya et al, 2018) were analyzed for DICER1 expression by **(F)** qPCR, **(G)** immunoblotting, and **(H)** immunohistochemistry. Actin was used as a loading control. **(I)** Cal62-luc cells were injected into the right thyroid lobe: a synthetic miR-146b inhibitor (Anti-146b) or a negative control was administered systemically via the retro-orbital vein. Left: endpoint (day 21) bioluminescent signal of the treated tumors. Right: tumor radiance quantification at the indicated time points in mice from treatment onset with the miRNA inhibitor (blue) or the negative control (green). **(J**) Representative immunohistochemistry with DICER1 antibody in orthotopic tumors. Values represent mean + SD (*n* = 3). **p< 0.01; ***p<0.001. For **(I),** values represent mean ± SEM. *p<0.05.

Given these data, we investigated the role of DICER1 in the aggressive traits induced by miR-146b overexpression, observing that its overexpression partly rescued the miR-146b-induced increase in proliferation, migration and invasion (Figure 1C-E). Overall, these results show that the most highly overexpressed miRs, and mainly miR-146-5p, target *DICER1* to mediate potential oncogenic effects and aggressiveness of thyroid cancer cells *in vitro*.

### Neutralization of miR-146b suppresses growth of established thyroid tumors *in vivo* and restores DICER1 expression

We next asked whether the continuous repression of DICER1 by endogenous miR-146b in aggressive thyroid cells is required for tumor growth *in vivo*. To test this, we first used a xenograft heterotopic model in which established solid tumors of Cal62 cells were treated intratumorally with a specific miR-146-5p oligonucleotide inhibitor (Anti-146b) or a control (Ramirez-Moya et al, 2018). Tumor growth was significantly blunted in the antimiR-treated tumors, as we demonstrated previously (Ramirez-Moya et al, 2018). We then analyzed DICER1 mRNA and protein levels in these xenograft tumors, finding that intratumoral anti-miR treatment rescued the levels of DICER1 (Figure 1F-H). This suggests that restoration of DICER1 expression by intratumoral inhibition of miR-146b contributes, at least in part, to tumor growth reduction. To better assess the therapeutic potential of antimiR-146b, we generated an orthotopic thyroid tumor in mouse. In this new model, luciferase-expressing cells (Cal62-luc) were directly injected into the right thyroid lobe of nude mice. After tumor establishment (3 weeks), 13 mice were randomized into two groups and were injected intravenously with antimiR-146b (*n*=8) or an appropriate control (*n*=5). Tumor volume was then followed for a further two weeks. Results showed that tumor growth was significantly delayed in the anti-miR-treated group (Figure 1I). Importantly, DICER1 expression in the primary thyroid tumor was higher after systemic antimiR-146b treatment (Figure 1J). Overall, these data reveal a functional pathway in thyroid tumors whereby inhibition of endogenous miR-146b expression, and therefore restoration of DICER1 expression, may be exploited therapeutically for thyroid cancer treatment.

### DICER1 plays a critical tumor suppressive role in thyroid cancer progression

As our data so far suggest an essential role for DICER1 in thyroid cancer, we first analyzed its protein expression levels in a panel of thyroid cancer cell lines. We found that DICER1 expression was lower in all human thyroid cancer cell lines examined than in control Nthy-ori cells (Supplementary Figure 2A-B). To examine the effects of DICER1 on tumor-specific phenotpyes *in vitro*, we performed loss-of-function and gain-of-function studies using the cell lines Cal62 and TPC1, which have intermediate levels of DICER1, and SW1736, which has low levels of DICER1(Supplementary Figure 2B).

We silenced DICER1 expression in Cal62 and TPC1 cells using a DICER1 siRNA (siDICER1). As expected, DICER1 silencing reduced the abundance of several mature miRs (miR-221-3p, miR-30a-5p, miR-21-5p, miR-146b-5p, miR-100-5p and miR-204-5p) (Figure 2A). Moreover, proliferation by PCNA immunoblotting, cell viability and DNA synthesis (Figures 2B–D), and also migration and invasion (Figure 2E, F) were all higher in DICER1-silenced Cal62 and TPC1 cells than in controls.

**Figure 2.**
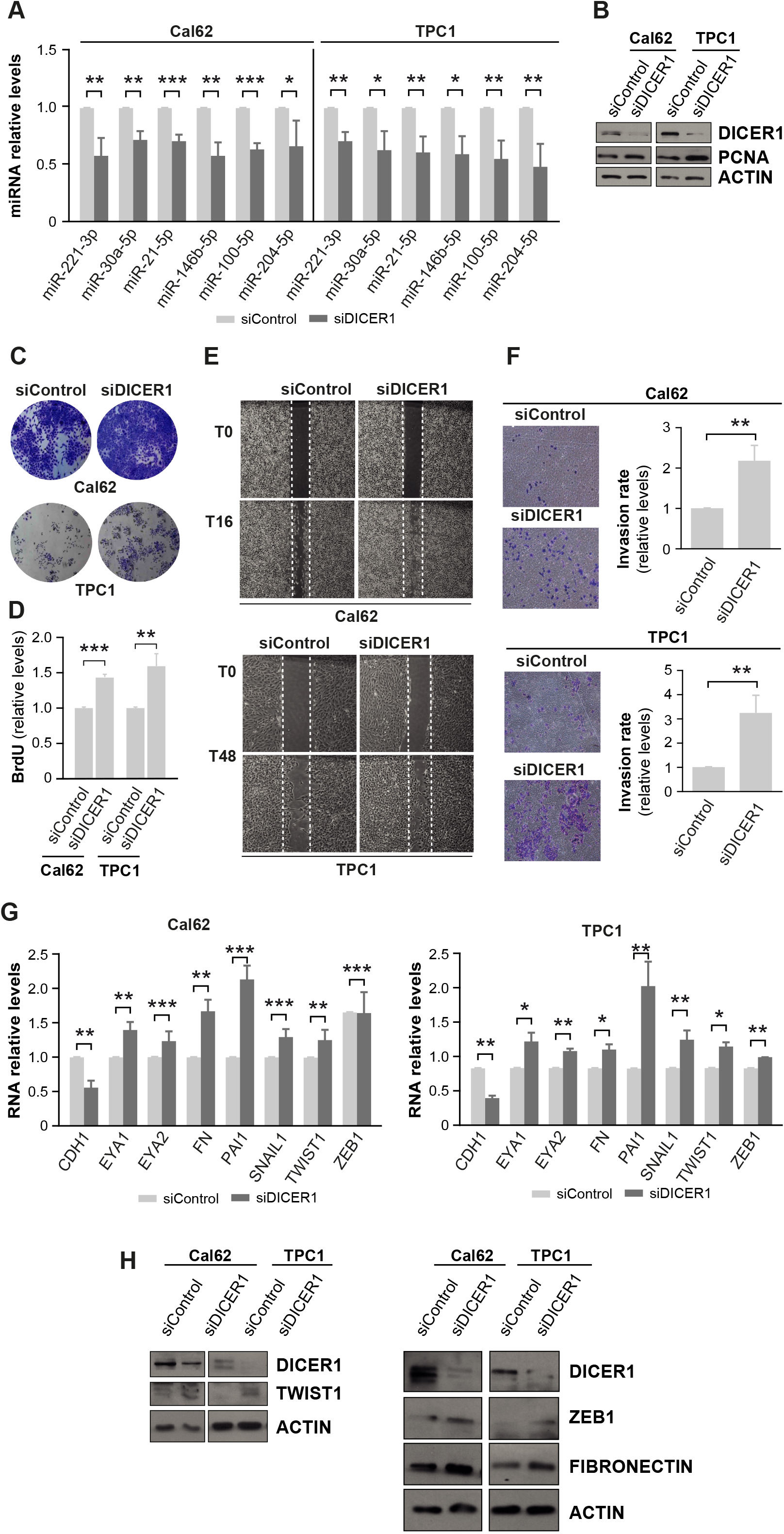
DICER1 silencing increases proliferation, migration and invasion *in vitro*. **(A–F)** Cal62 and TPC1 cell lines were transfected with a siRNA against DICER1 (siDICER1) or a control siRNA (siControl) and analysis was performed 48 h later. **(A)** Relative expression levels of miRs in Cal62 and TPC1 cells. **(B)** Immunoblot of DICER1 and proliferating cell nuclear antigen (PCNA). **(C)** Representative images of crystal violet-stained Cal62 and TPC1 cells. **(D)** BrdU incorporation relative to siControl-transfected cells. **(E)** Representative images from a wound healing assay at 0 and 16 h (Cal62, left) or 0 and 48 h (TPC1, right) after scratching. **(F)** Quantification of the invasion rates. Left: representative images of the lower chamber (invading cells). Right: cell invasion rates relative to siControl cells. (**G** and **H**) Cal62 and TPC1 cell lines were transfected with an siRNA against DICER1 (siDICER1) or a control siRNA (siControl). **(G)** Relative mRNA levels of EMT genes *CDH1*, *EYA1*, *EYA2*, *FN*, *PAI1*, *SNAIL1*, *TWIST1*, and *ZEB1* 48 h after siRNA transfection by qRT-PCR. **(H)** Immunoblot for DICER1 and TWIST1 (left) and DICER1, ZEB1 and FN (right) 48 h after siRNA transfection. Values represent mean + SD (*n* = 3). *p<0.05; **p<0.01; ***p<0.001.

We also examined whether the expression of protein markers involved in epithelial-mesenchymal transition (EMT) was altered in DICER1-silenced thyroid cells. Whereas DICER1 silencing significantly decreased the mRNA levels of the epithelial protein E-cadherin (CDH1), it increased the expression of several mesenchymal genes/proteins, including EYA1, EYA2, fibronectin, PAI1, SNAIL1, TWIST1 and ZEB1, in both cell lines (Figure 2G, H). These data demonstrate that DICER1 is a critical player in thyroid tumor progression, as its knockdown copied the phenotype observed by endogenous miR-146b overexpression in an orthogonal fashion, and that DICER1 may play a role in EMT and metastasis formation in PTC.

In contrast to the loss-of-function studies above, SW1736 cells ectopically overexpressing DICER1 showed an expected induction of several DICER-dependent mature miRs (Supplementary Figure 3A). Moreover, proliferation, migration and invasion were all significantly decreased in cells overexpressing DICER1, together with the corresponding changes in the expression of EMT markers (Supplementary Figure 3B–G). Overall, these data unequivocally establish a novel role of DICER1 as a tumor suppressor in thyroid follicular cells.

We next explored which signaling pathways were involved in the oncogenic effect mediated by DICER1 repression. MAPK and PI3K signaling pathways are essential in thyroid carcinogenesis (Riesco-Eizaguirre & Santisteban, 2016). We observed that PI3K activity, evaluated by AKT phosphorylation, was upregulated in DICER1-silenced Cal62 and TPC1 cells (Supplementary Figure 4A). By contrast, MAPK activity, evaluated by ERK phosphorylation, was similar irrespective of DICER1 expression levels (Supplementary Figure 4B). These data confirm that DICER1 downregulation contributes to PI3K signaling hyperactivation and point to its important role in the transition to a more aggressive state in thyroid cancer.

### Association between DICER1 expression and worse clinical outcome in thyroid cancer patients

To study the clinical relevance of DICER1 repression in thyroid cancer, we compared the mRNA levels of DICER1 in tumor samples and normal tissues using data acquired from TCGA FireBrowse portal and the Morpheus tool. We observed that DICER1 levels were decreased in several cancer types (Figure 3A), and thyroid cancer had the third greatest change in DICER1 expression overall when compared with normal tissue (Figure 3B) (normal tissue *n*=59, tumor samples *n*=501). Of note, thyroid metastases (*n*=8) exhibited even lower levels of *DICER1* that primary tumor samples or normal tissue (Figure 3B), suggesting a role for DICER1 downregulation in tumor progression. To extend these observations, we surveyed DICER1 mRNA levels in an independent paired cohort of 7 PTC patients, finding that levels were lower in tumor samples than in contralateral normal thyroid tissue in most patients (Figure 3C, left). Globally, we observed a significant decrease in DICER1 mRNA levels of ~46% (Figure 3C, right).

**Figure 3.**
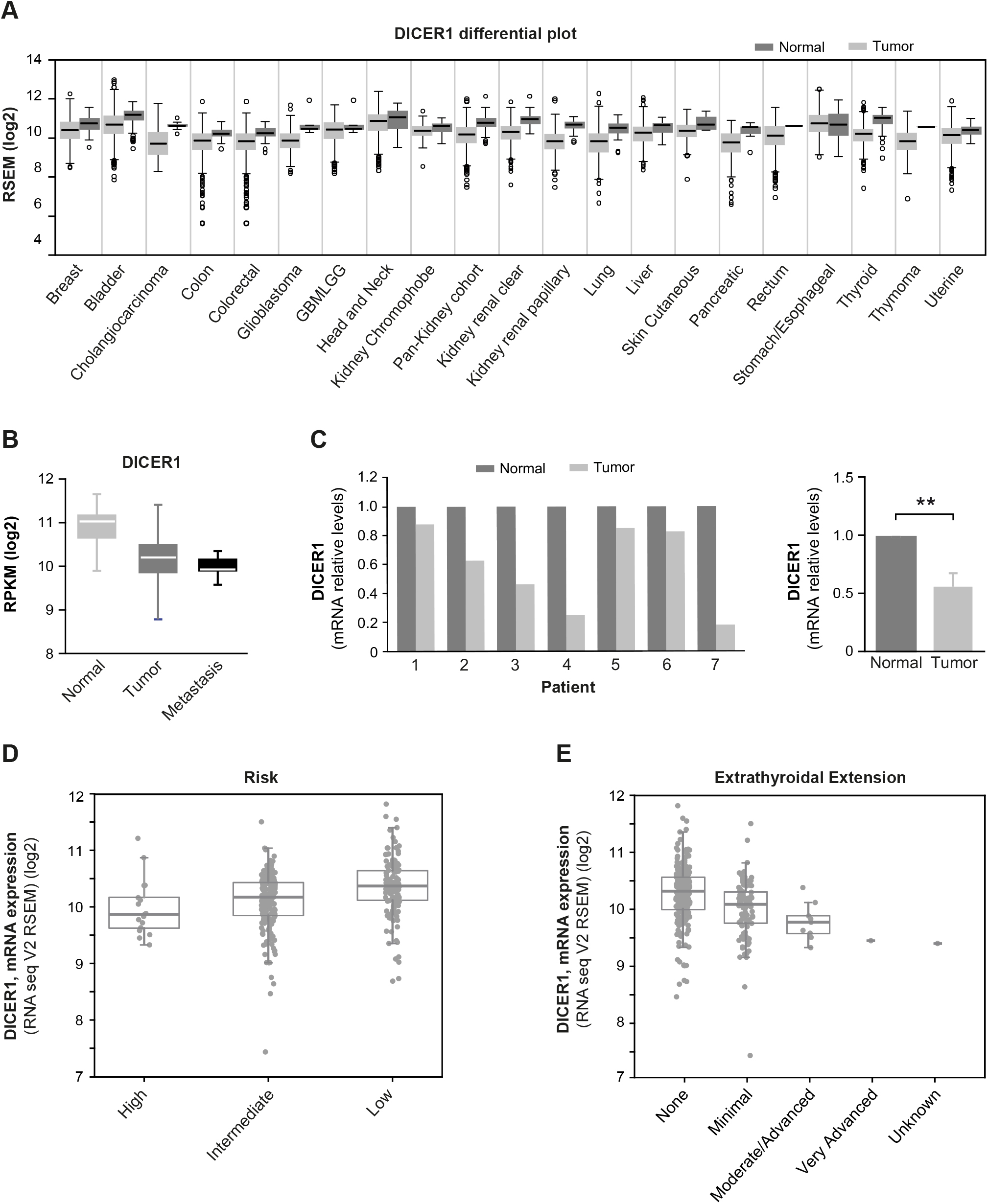
DICER1 is downregulated in human PTC tumors. **(A)** DICER1 mRNA expression levels in the indicated cancer types obtained by Firebrowse analysis of the TCGA database. **(B)** Box plot of DICER1 mRNA expression levels in thyroid normal tissue, PTC, and metastases: data was obtained from the TCGA database. **(C)** Left: relative DICER1 mRNA levels in 7 PTC patients (contralateral and normal thyroid tissue). Right: total average of DICER1 mRNA relative levels. **(D** and **E)** Correlation between DICER1 mRNA levels in high, intermediate or low risk **(D),** or extrathyroid extension **(E)** by analyzing the cBioPortal, where the TCGA datasets are deposited. Values represent mean + SEM. **p<0.01

To further explore the implications of DICER1 dysregulation in thyroid cancer, we sought to study its correlation with aggressive clinical features. TCGA data from the public databases cBioportal and Cancer Regulome indicated that DICER1 levels were inversely correlated with the risk of recurrence (Figure 3D) and with extrathyroidal extension (Figure 3E). Moreover, DICER1 was more downregulated in tumors classified within the miR clusters 5 and 6, described in TCGA as the less-differentiated tumors with high risk of recurrence (Supplementary Figure 5A) ( Cancer Genome Atlas Research Network, 2014; Riesco-Eizaguirre & Santisteban, 2016). Overall, these data strongly suggest an important role for DICER in thyroid tumor progression.

Because the maturation of most miRNAs is blocked in the absence of DICER1 (Bernstein et al, 2003), we analyzed the global expression of the miRs differentially expressed in thyroid cancer. Through the analysis of TCGA data, we found 93 miRs differentially expressed between samples of PTC and normal tissue: whereas 20 miRs were upregulated, as many as 73 were downregulated approximately ~3.5-fold or more, false discovery rate [FDR] <0.01 and fold change (FC) >2 for upregulated miRs and FC <0.75 for downregulated miRs) (Supplementary Figure 5B). Similar results were obtained when we analyzed the data obtained from an independent cohort of patients published by the group of De la Chapelle (Swierniak et al, 2013) (results not shown). The higher number of downregulated miRs relative to upregulated miRs in thyroid tumors suggests an important role of potential tumor suppressive miRs in this cancer type, and are in line with the findings of DICER1 downregulation.

### Individual downregulated miRs rescue DICER1-silencing effects

Since we clearly established a direct role of DICER1 repression by miR-146-5p and potentially other miRs in PTC, and given its tumor suppressive effect, we next sought to address further downstream DICER1 effector miRs relevant to this phenotype. In this regard, we and others recently profiled repressed miRs in PTC with potential tumor suppressive functions ( Cancer Genome Atlas Research Network, 2014; Riesco-Eizaguirre et al, 2015), which included miR-30a and miR-100. These miRs are reported to be essential for EMT and the mesenchymal phenotype by, for example, targeting lysyl oxidase and promoting SNAIL2 expression (Boufraqech et al, 2015; Hebrant et al, 2014; Lima et al, 2017) or have been described as tumor suppressors in other cancer entities. To assess whether the effects of DICER1 silencing are directly attributed to these repressed miRs, we restored miR-30a and miR-100 expression in DICER1-silenced cells and monitored the resulting phenotype. Indeed, transfection of miR-30a or miR-100 reversed the DICER1-silenced phenotypes, and reduced proliferation (Figure 4A, B), migration (Figure 4C) and invasion (Figure 4D). These results indicate that inhibition of miRs with tumor-suppressor functions, such as the DICER effectors miR-30a or miR-100, is critical for the proliferative and motile properties of malignant tumoral thyroid cells.

**Figure 4.**
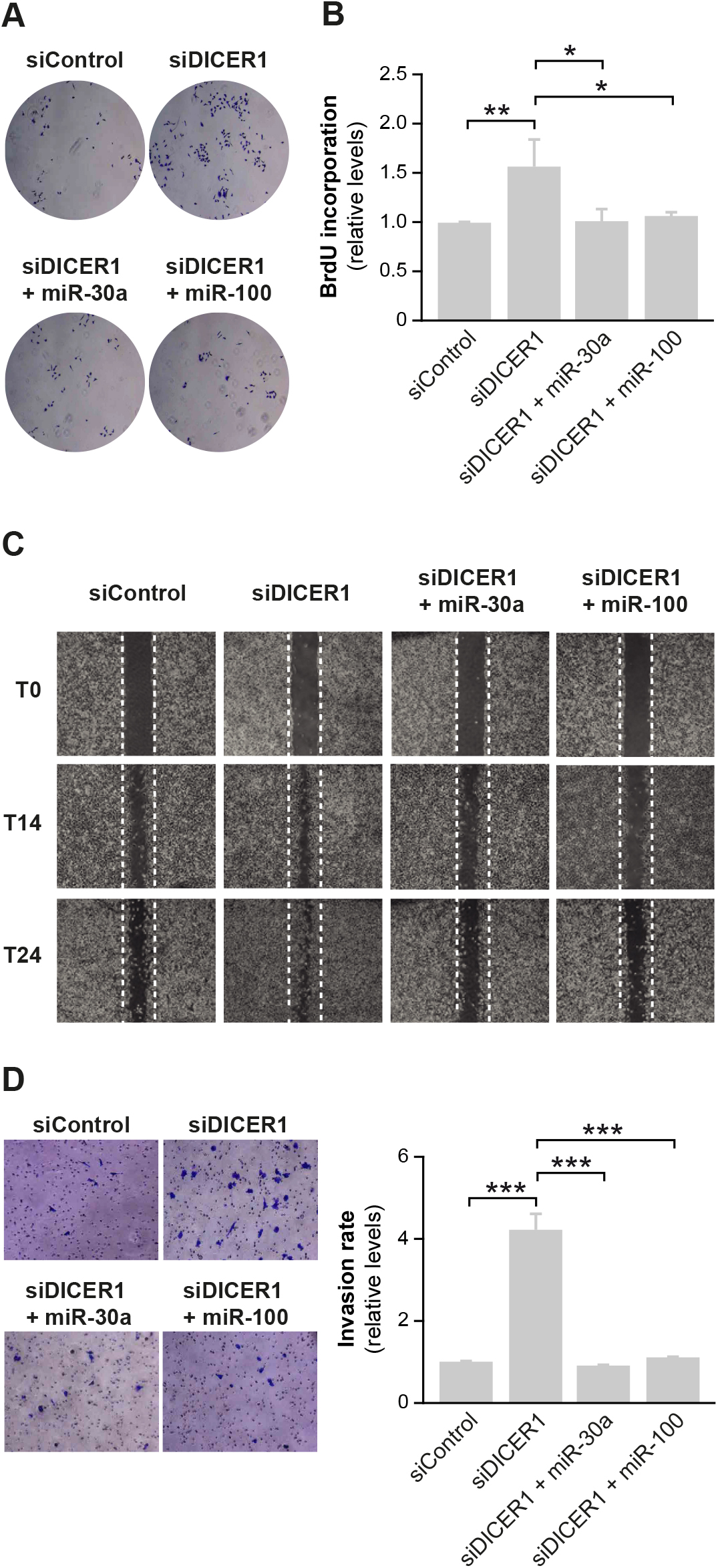
Downregulated miR-30a and miR-100 in thyroid cancer block DICER1 silencing-induced effects. Cal62 cells were silenced for DICER1 (siDICER1) and co-transfected with miR-30a or miR-100 plasmids. Control cells were transfected with siControl and the empty DICER1 vector. **(A)** Representative images of cells transfected with siControl, siDICER1, siDICER1 + miR-30a, and siDICER + miR-100 at 48 h post-transfection and stained with crystal violet. **(B)** BrdU incorporation relative to cells transfected with the control siRNA (siControl). **(C)** Representative images from a wound healing assay 0, 14 and 24 h after scratching in the above-mentioned conditions. **(D)** Quanatification of invasion capacity was analyzed in Matrigel-coated transwells. Left: representative images of the lower chamber (invading cells). Right: cell invasion relative to siControl cells. Values represent mean + SD (n = 3). *p<0.05; **p<0.01; ***p<0.001.

### Induction of miR processing with the small molecule enoxacin decreases cell aggressiveness *in vitro* and tumor growth *in vivo*

Although many studies have investigated the impact of miRs on cancer biology, miR-based cancer therapy is still in its early stages and is mostly limited to targeting a single miR (Adams et al, 2017; Gilles et al, 2018). However, because global down-regulation of miR expression and corresponding DICER1 repression appears to play a role in thyroid cancer, restoration of their levels or enhancing miR biogenesis might represent an attractive approach in cancer therapy. Against this background, we treated Cal62, TPC1 and SW1736 cells with the small molecule enoxacin, which enhances miR maturation by binding to TRBP. Consistent with its activity against other types of tumors (Melo et al, 2011), the enhancement of DICER1 complex enzymatic activity by enoxacin resulted in an increase in the expression of all tested mature miRs, including tumor suppressor miRs (Figure 5A). Thus, we next investigated whether enoxacin could act as a cancer growth inhibitor by examining its effects in thyroid cancer cell lines. Cal62, TPC1 and SW1736 cells treated with enoxacin showed a dramatic decrease in proliferation, cell migration and cell invasion capacity together with a decrease of PCNA protein expression levels (Figure 5B-F). Furthermore, inhibition of migration and invasion by enoxacin was accompanied by a decrease in the expression of the EMT markers fibronectin, N-cadherin, ZEB and TWIST (Figure 5G). Finally, enoxacin treatment also decreased pAKT expression to almost undetectable levels in the three thyroid cancer cell lines tested (Figure 5H).

**Figure 5.**
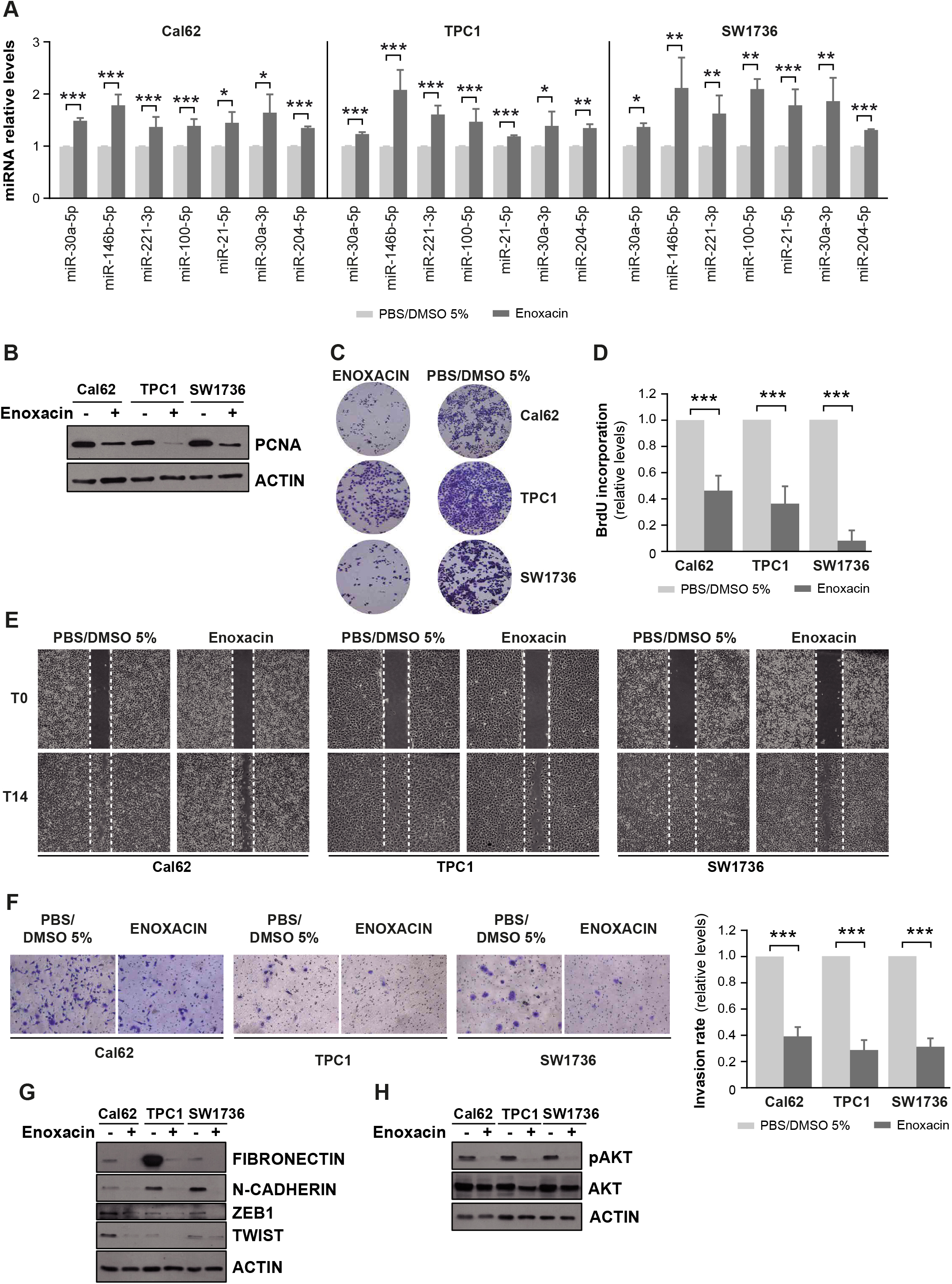
Enoxacin induces miR expression and decreases cell proliferation, migration and invasion *in vitro*. Cal62, TPC1 and SW1736 cell lines where treated with enoxacin or PBS + 5% DMSO (control) for 5 days. **(A)** Relative expression levels of miR-30a-5p, miR-146b-5p, miR-221-3p, miR-100-5p, miR-21-5p, miR-30a-3p, and miR-204-5p in cells. **(B)** Immunoblot od proliferating cell nuclear antigen (PCNA) and actin, as a loading control. **(C)** Representative images of cells treated as described and stained with crystal violet. **(D)** BrdU incorporation of cells treated with enoxacin relative to control-treated cells. **(E)** Representative images from a wound healing assay 0 and 14 h after scratching. **(F)** Quantification of invasion rates. Left: representative images of the lower chamber (invading cells). Right: cell invasion rates relative to control-treated cells. **(G)** Immunoblot for fibronectin, N-cadherin, ZEB1 and TWIST1 and, **(H)** pAKT and AKT using actin as a loading control. Values represent mean + SD (*n* = 3). *p<0.05; **p<0.01; ***p<0.001.

Finally, to translate these *in vitro* findings to an *in vivo* thyroid cancer model, we tested enoxacin in the orthotopic mouse model of human thyroid cancer. Accordingly, Cal62-luc cells where injected into the right thyroid lobe to establish tumors as described above, and 15 mice were randomized into two groups: a test group (*n*=8) treated daily by intraperitoneal injection of enoxacin (15 mg/kg) over 28 days, and a vehicle control group (*n*=7) treated with 5% DMSO in phosphate buffered saline (PBS). Tumor bioluminiscent signals were analyzed *in vivo* twice weekly to calculate tumor growth. Compared with the control group, we found that enoxacin significantly lessened the radiance increase (Figure 6A left) and decreased tumor growth (Figure 6A right). We confirmed the upregulated expression of critical miRs in tumors from enoxacin-treated mice (Figure 6B), and the downregulation of the EMT markers fibronectin and N-cadherin and the proliferation marker PCNA (Figure 6C). Overall, these data suggest that enoxacin enhances miR production and, as a consequence, re-establishes DICER1 miRNA processing, eliciting its antitumor effect. We thus conclude that the miR biogenesis machinery plays a critical role in maintaining a differentiated thyroid state and its impairment induces an aggressive tumoral state and small molecule treatment with enoxacin is effective against PTC tumor progression in *in vivo* mouse models.

**Figure 6.**
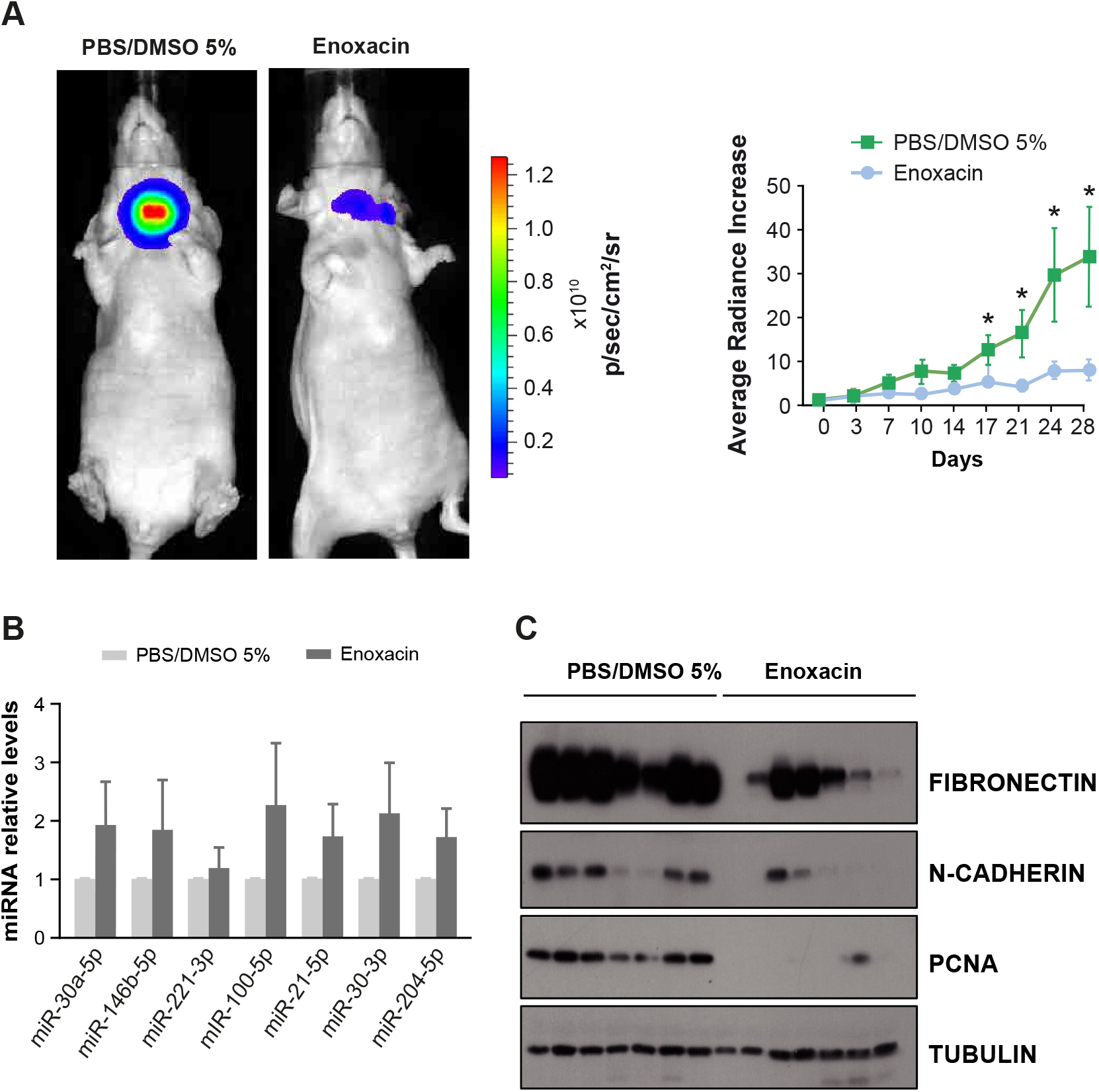
Enoxacin impairs established human orthotopic thyroid tumor growth and decreases the expression of EMT-related genes *in vivo*. **(A–C)** Cal62-luc cells were injected into the right thyroid lobe. After the orthotopic model was established, enoxacin or vehicle was administered intraperitoneally. **(A)** Left: the image shows the endpoint (day 28) bioluminescent signal of the tumors imaged with the IVIS-Lumina II Imaging System. Right: tumor radiance quantification at the indicated time points in mice from treatment onset with enoxacin (blue) or vehicle (green). **(B)** RNA was obtained from the tumors and miR expression levels were determined. Shown are the relative intratumoral levels of miR-30a-5p, miR-146b-5p, miR-221-3p, mIR-100-5p, miR-21-5p, miR-30-5p, and miR-204-5p. **(C)** Immunoblot of intra-tumoral expression of fibronectin, N-cadherin and PCNA in enoxacin or vehicle-treated mice. Values represent mean ± SEM. *p<0.05.

## Discussion

Here, we provide evidence that thyroid cancer cells use global downregulation of an miR sub-network to enhance disease aggressiveness, including the induction of proliferation and metastatic capacity. Global downregulation of miR maturation has been described in many other tumor types (Lin & Gregory, 2015; Lu et al, 2005; Martello et al, 2010; Ozen et al, 2008; Thomson et al, 2006) including thyroid cancer, where it has been suggested that distinct miR expression patterns can distinguish ATC from FTC or from PTC (Braun et al, 2010). Our results give direct support for and considerably extend this concept by identifying a mechanism by which miR down-modulation leads to thyroid tumor progression. We show that the most overexpressed miRs in thyroid cancer (miR-146b, - 21, −222, −221, −182) commonly target DICER1, the RNAse III-type enzyme that controls functional maturation of most miRs, inhibiting its expression as illustrated in Figure 7. These results are reinforced by the exhaustive analysis of TCGA data (Cancer Genome Atlas Research Network, 2014) that we carried out through different bioinformatics analyses. Among the miRs enriched in thyroid cancer, we focused our attention on miR-146b, as we have recently characterized its oncogenic activity in thyroid cancer (Ramirez-Moya et al, 2018). The miR-146b-DICER1 interaction revealed in this work has profound consequences for the induction of an aggressive phenotype, as ectopic DICER1 expression or enzymatic activation by enoxacin rescues the aggressive phenotype induced by miR-146b. Since miR-146b is both produced by and regulates DICER1, this mutual feedback relationship allows for a low level of DICER1 in thyroid cancer but not its complete loss and thus sufficient DICER1 levels are maintained for cell survival and growth.

**Figure 7.**
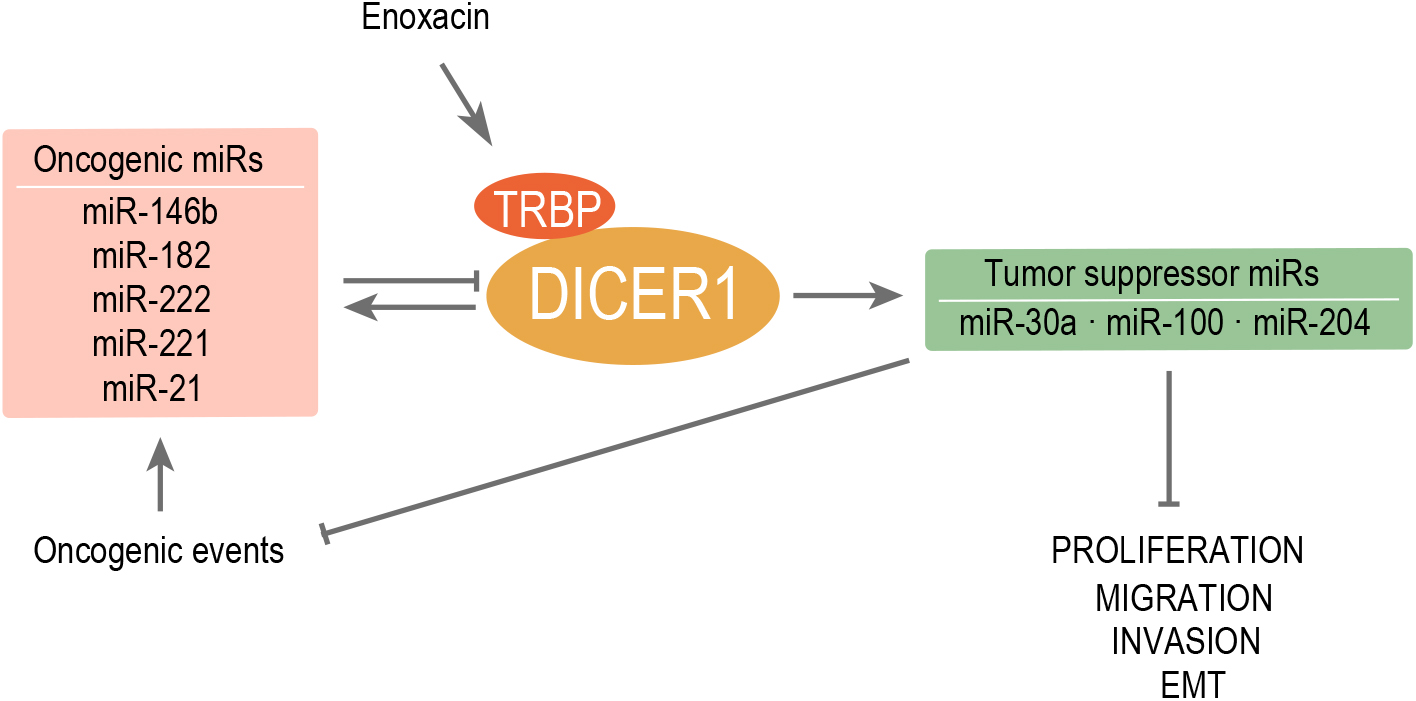
Summary of established DICER1-miRNA regulatory network in thyroid cancer. Overexpressed (oncogenic) miRs in thyroid cancer directly target DICER1. Also, DICER1 upregulates tumor suppressor miRs, which in turn inhibit proliferation, migration, invasion and EMT. This effect can be phenocopied when increasing the DICER1 complex activity with the small molecule enoxacin The arrows denote induction and the truncated lines repression. Dashed line denotes bioinformatic predictions

We show that the reduction of DICER1 expression by miR-146b is critical for the execution of a metastatic program. Accordingly, DICER1 overexpression should phenocopy the loss of miR-146b and oppose metastasis. Repression of DICER1 is one, but not the only oncogenic action of miR-146b, as we have demonstrated that it also represses the tumor suppressor PTEN, leading to an activation of the oncogenic PI3K/AKT pathway (Ramirez-Moya et al, 2018). In accordance with this is the finding that DICER1 silencing increases AKT activation, which reinforces the importance of this pathway in thyroid tumorigenesis. We believe that activation of PI3K/AKT together with the global downregulation of miRs could contribute to a more aggressive tumor phenotype. In our *in vivo* studies, antimiR-146b treatment suppressed tumor growth and reversed DICER1 expression. These findings have implications for the treatment of thyroid cancer, as the positive effects of antimiR-146b in our experimental models suggest that modulation of miR-146b by RNA-based therapeutics may prove clinically useful.

Conceptually, some of the present data may seem paradoxical, as DICER1 is downregulated in thyroid cancer, but some miRs are upregulated. However, it is known that, among other phenomena, the activation of oncogenes causes upregulation of specific miRs (Kumar et al, 2007). For example, conditional activation of BRAFV600E and RET/PTC3 in normal thyroid cells induces the expression of miR-146b (Geraldo et al, 2012), which can explain the co-existence of a general miR downregulation in tumors with low DICER1 levels and the upregulation of miR-146b. In addition, other oncogenic pathways in thyroid cancer have been described to increase miR-146b levels such as NF-κB (Pacifico & Leonardi, 2010; Taganov et al, 2006) or PI3K/AKT pathways (Ramirez-Moya et al, 2018). Another possible explanation for miR upregulation could be gene demethylation. Consistent with this is the result of our analysis of TCGA database, showing that miR-146b is demethylated in thyroid cancer and its metastasis (Supplementary Figure 6A), whereas downregulated miRs have almost no change in methylation levels (not shown).

Concerning the consequences of loss of DICER1 expression in more aggressive phenotypes, analysis of TGCA data showed an association between low DICER1 expression and thyroid metastasis, high risk, and extrathyroidal extensions. Indeed, we confirmed low DICER1 mRNA levels in a panel of seven PTC thyroid tumors. Taken together, these data suggest that, similar to other tumor types (Lin & Gregory, 2015; Su et al, 2010; Yan et al, 2012), low DICER1 levels in thyroid cancer are associated with advanced tumor stage and poor clinical outcome, an observation that has been previously reported (Erler et al, 2014). Moreover, the present work is the first to provide functional evidence that DICER1 acts as a strong tumor suppressor in thyroid cancer. Although rare, several DICER1 mutations, both germinal and somatic, have been described in human thyroid cancers (Khan et al, 2017; Rutter et al, 2016), resulting in a loss of function. These studies underscore the pathophysiological relevance of the miR biogenesis machinery in thyroid tumors.

Our observations in human thyroid cancer were supported by cellular and in mouse models, allowing us to study the functional consequences of DICER1 downregulation. In loss-of-function experiments, we observed a downregulation of miRs in DICER1-silenced cells, which induced cell proliferation, migration and invasion and also the EMT phenotype characteristic of metastatic capacity (Figure 7). The opposite results were obtained in gain-of-function experiments, validating our experimental approach. Regarding proliferation, our data contrast with a report showing that DICER1 overexpression positively regulates thyroid cell proliferation (Penha et al, 2017). However, another group has shown that the knockout of *Dicer1* in the thyroid of mice resulted in a marked increase of proliferation (Rodriguez et al, 2012). In addition, the extensive data derived from TCGA analysis and all the novel data reported here strongly support the concept that DICER1 inhibits cell proliferation and consequently acts as a tumor suppressor and not an oncogene. The crucial role of DICER1 was confirmed by the finding that total abolition of DICER1 expression by CRISPR/Cas9 gives rise to non-viable cells (Supplementary Figure 6B).

We have demonstrated that global miR downregulation induced by DICER1 promotes thyroid cell aggressiveness. Noteworthy, thyroid cancer progression leads to less upregulated and more downregulated miRs (Hebrant et al, 2014; Saiselet et al, 2016). This observation is consistent with the hypothesis that the biogenesis machinery breaks down as the tumor moves towards a more aggressive stage. That an aggressive tumoral behavior becomes manifest after the general downregulation of miRs points to an important contribution of this process to the aggressive phenotype, including the downregulation of tumor suppressor miRs. This is supported by our data in DICER1-silenced cells where the expression of tumor suppressors miR-30a and miR-100 recover the normal thyroid phenotype. Thus, further study of downregulated tumor suppressor miRs that are essential to protect cells from activation of oncogenes could be an important approach against cancer.

It is important to find strategies to restore global miR expression as a potential therapy in thyroid tumors. Regarding the restoration of miRs with tumor suppressor roles, fewer examples exist, but it is reasonable to think that restoring global miR levels could have a therapeutic effect. Along this line, we used the small molecule enoxacin, which binds and induces TRBP, a component of the DICER1 complex (Melo et al, 2011). Enoxacin enhanced the production of miRs and decreased cell aggressiveness *in vitro* and tumor growth *in vivo*. We not only show that this molecule has a cancer-specific inhibitory effect, but we also provide evidence that activating miR biogenesis is an interesting step towards the potential application of miR-based therapy for the treatment of thyroid cancer. This therapy is promising in other pathologies, as shown in mouse models of amyotrophic lateral sclerosis, where it has been suggested that DICER1 and miRs affect neuronal integrity and are possible therapeutic targets (Emde et al, 2015).

In summary, our work reveals some key aspects on the regulation and function of DICER1 in thyroid cancer. Paradoxically, DICER1 is downregulated in thyroid cancer through the upregulation of targeting miRs, particularly miR-146b. DICER1 repression functionally leads to global downregulation of the miR network, inducing increased malignancy. Our data may have important clinical implications as DICER1 may convey prognostic information and, more importantly, point to a potential therapeutic approach based on the restoration of global miR levels *via* the DICER1 pathway or using DICER1-targeting miR inhibitors in patients with thyroid and other cancers.

## Material and Methods

### Bioinformatic predictions

The TCGA database was queried to assess correlations between mRNA and miRNA levels and clinical features. Firebrowse (http://firebrowse.org) was used to analyze *DICER1* mRNA levels in different tumor types. The cBioPortal (http://www.cbioportal.org) and the Cancer Regulome Explorer (http://explorer.cancerregulome.org) data portals were used to obtain the correlations using the thyroid carcinoma dataset (THCA). The miRanda algorithm (http://www.microrna.org) was used to predict hypothetical associations between microRNAs and mRNAs.

### Patients

Samples of PTC tumors and contralateral normal thyroid tissue from the same patients (*n*=7) were collected at the Biobank of the Hospital Universitario La Paz (Madrid, Spain). Written informed consent was obtained previously from all the patients in accordance with the protocols approved by the Ethic Committee.

### RNA quantification

RNA was extracted using TRIzol (Invitrogen). RT-PCR and qRT-PCR were performed as described in Supplementary Materials and Methods.

### Cell culture and transfection

The human thyroid cancer cell lines of PTC (BCPAP and TPC1) and ATC (Ocut2, Ktc2, Cal62, T235, Hth83, Hth74, and SW1736) were cultured as described (Sastre-Perona et al, 2016). Rat PCC13 cells (Fusco et al, 1987) and the human primary thyroid follicular cell line Nthy-ori 3-1 (Ramirez-Moya et al, 2018), derived from normal thyroid tissue, were cultured as described. Stable cell lines, transfections and Luciferase activity were performed as described in Supplementary Materials and Methods.

### Plasmids, constructs, and siRNA

The *DICER1* expression vector was kindly provided by Dr Richard Gregory (Boston Children’s Hospital, Boston) (Chendrimada et al, 2005). The pGL3-DICER-Prom vector was a gift from Dr. David Fisher (Addgene plasmid #25851) (Levy et al, 2010) and the pIS-DICER1 long UTR was a gift from Dr David Bartel (Addgene plasmid #21649) (Mayr & Bartel, 2009). The luciferase and GFP-expressing vector, CMV-Firefly luc-IRES-EGFP, was constructed by Dr. J. Bravo (IQAC-CSIC), and the Cal62 human tumoral thyroid cell line stably expressing this vector (Cal62-Luc) was generated by Dr. Eugenia Mato (IB, Sant Pau).

The pre-miR-146b construct was previously cloned into a pEGP expression vector (Cell Biolabs) (Ramirez-Moya et al, 2018). Pre-miR-30a and pre-miR-100 miRs were amplified from genomic DNA and cloned into the pEGP-miR expression vector. DICER1 siRNA was purchased from Thermo Fisher (Silencer^®^ Select Pre-Designed siRNA Dicer1, s23754).

To produce DICER1 knock-out cell lines, we used the CRISPR/Cas9 technique as described in Supplementary Materials and Methods.

### Protein extraction, western blotting and immunohistochemistry

Cells were lysed and proteins extracted as described (Ramirez-Moya et al, 2018). Western blotting and inmunohistochemestry procedures are detailed in Supplementary Materials and Methods.

### Cell proliferation assays

Proliferation was determined by proliferating cell nuclear antigen (PCNA) expression, crystal violet staining and bromodeoxyuridine (BrdU) incorporation, and measured as described (Ramirez-Moya et al, 2018; Sastre-Perona et al, 2016). Full details are in Supplementary Materials and Methods.

### Migration and invasion assays

Wound healing and cell invasion assays were performed as described (Ramirez-Moya et al, 2018), and as detailed in Supplementary Materials and Methods.

### Enoxacin treatment in vitro

To stimulate miR expression, cells were treated with enoxacin, sodium salt (EMD Millipore Corp.; cat. no. 557305) at 40 μg/mL diluted in PBS with 5% DMSO, for 5 days (Melo et al, 2011).

### In vivo studies

Animal experimentation was performed in compliance with the European Community Law (86/609/EEC) and the Spanish law (R.D. 1201/2005), with the approval of the Ethics Committee of the Consejo Superior de Investigaciones Científicas (CSIC, Spain).

Xenograft models have been previously described (Ramirez-Moya et al, 2018). Orthotopic implantation was performed as described in Supplementary Materials and Methods.

### Statistical analysis

Results are expressed as the mean ± SD of at least three different experiments performed in triplicate. Results from the *in vivo studies* and patient analysis are expressed as the mean ± SEM. Statistical significance was determined by Student’s t-test analysis (two-tailed) and differences were considered significant at a P value <0.05.

## Author Contributions

J.R-M conducted the experiments. J.R-M and L.W-L performed the *in vivo* experiments. J.R-M acquired, analyzed the data and conducted bioinformatic analysis. J.R-M, L.W-L and P.S designed research studies. G. R-E provided thyroid tumors samples and clinical information of the patients. J.R-M, and P.S wrote de manuscript and G.R-E contribute to critical discussion.

## The paper explained

### Problem

The global downregulation of miRNAs is emerging as a common hallmark of cancer, however the underlying mechanisms and the phenotypic advantages provided to cells by reduced miR expression is not well known.

### Results

In this study, we took advantage of the genomic analysis of thyroid cancer, where the well-differentiated tumors present both up- and down-regulated miRs, whereas de-differentiated and aggressive one show almost exclusively down-regulated miRs. Analyzing human thyroid tumors, *in vitro* culture and thyroid tumor mouse models, we demonstrate that the main upregulated miR-146b suppress DICER1 expression, an essential enzyme for processing miR precursors to mature miRs. We have defined an association between low DICER1 levels and a worse clinical outcome, providing information that DICER1 acts as a tumor suppressor. The rescue of DICER1 expression or activity using anti-miR or the small molecule enoxacin, reduced thyroid cancer aggressiveness both *in vitro* and *in vivo*.

### Impact

This study highlights a potential therapeutic application of RNA-based therapies including miR-inhibitors and restoration of the biogenesis machinery, which may provide treatments for thyroid cancer and other cancers.

## Acknowledgements

We are grateful to Dr. Richard I Gregory (Stem Cell Program, Boston Children’s Hospital, Boston) and Jochen Imig (Department of Pathology, NYU School of Medicine, New York) for their excellent suggestions throughout the course of this work. We thank Dr. Kenneth McCreath for helpful comments on the manuscript. We acknowledge Javier Perales-Patón (CNIO, Madrid, Spain) for his advice in the bioinformatics analysis, Andrea Martinez-Cano for technical assistance and Javier Perez for artwork. We also thank the Histology Facility at CNB-CSIC for the histological preparation of biological samples.

